# Empagliflozin inhibits proximal tubule NHE3 activity, preserves GFR and restores euvolemia in nondiabetic rats with induced heart failure

**DOI:** 10.1101/2020.07.16.207118

**Authors:** Flávio A. Borges-Júnior, Danúbia Silva dos Santos, Acaris Benetti, Renato O. Crajoinas, Ednei L. Antonio, Leonardo Jensen, Bruno Caramelli, Gerhard Malnic, Paulo J. Tucci, Adriana C. Girardi

## Abstract

**Background:** Sodium-glucose cotransporter type 2 (SGLT2) inhibitors reduce the risk of heart failure (HF) mortality and morbidity, regardless of the presence or absence of diabetes, but the mechanisms underlying this benefit remain unclear. We tested the hypothesis that the SGLT2 inhibitor empagliflozin inhibits proximal tubule (PT) Na^+^/H^+^ exchanger isoform 3 (NHE3) activity and improves renal salt and water handling in nondiabetic rats with HF.

**Methods and Results:** Male Wistar rats were subjected to myocardial infarction or sham operation. After four weeks, rats that developed HF and sham rats were treated with empagliflozin (EMPA) or untreated for an additional four weeks. EMPA-treated HF rats displayed lower levels of serum brain natriuretic peptide (BNP) and lower right ventricle and lung weight to tibia length than untreated HF rats. Upon saline challenge, the diuretic and natriuretic responses of EMPA-treated HF rats were similar to those of sham rats and were higher than those of untreated HF rats. Additionally, EMPA treatment normalized the glomerular filtration rate and proteinuria in HF rats. PT NHE3 activity was higher in HF rats than in sham rats, whereas treatment with EMPA markedly reduced NHE3 activity. Unexpectedly, SGLT2 function and protein and mRNA abundance were upregulated in the PT of HF rats.

**Conclusion:** Collectively, our data show that the prevention of HF progression by empagliflozin is associated with inhibition of PT NHE3 activity and restoration of euvolemia. Moreover, we propose that the dysregulation of PT SGLT2 may be involved in the pathophysiology of nondiabetic HF.

**SIGNIFICANCE STATEMENT:** SGLT2 inhibitors represent a class of drugs that were originally developed for improving glycemic control. Cardiovascular outcome trials that were designed to evaluate cardiovascular safety yielded unexpected and unprecedented evidence of the cardiorenal benefits of SGLT2 inhibitor. Many hypotheses have been proposed to explain the mechanisms underlying these effects. Our study demonstrates that SGLT2 inhibition restores extracellular volume homeostasis in nondiabetic heart failure (HF) rats by preserving GFR and inhibiting proximal tubule NHE3-mediated sodium reabsorption. The attenuation of kidney dysfunction may constitute an essential mechanism by which SGLT2 inhibitors attenuate HF development and progression either in the presence or absence of diabetes.

## INTRODUCTION

Sodium-glucose cotransporter type 2 (SGLT2) inhibitors, also known as gliflozins, suppress glucose reabsorption in the renal proximal tubule (PT), leading to substantial glycosuria, thereby lowering hyperglycemia in patients with type 2 diabetes (T2D)^1,2^ Gliflozins were initially developed as antidiabetic agents but have recently emerged as among the most impactful cardiovascular (CV) drugs. Three cardiovascular outcome trials consistently showed that SGLT2 inhibitors remarkably reduce cardiovascular mortality and hospitalization for heart failure (HF) in T2D patients^3–5^. Most recently, the phase III DAPA-HF (Dapagliflozin and Prevention of Adverse-outcomes in Heart Failure)^6^ study has announced that dapagliflozin reduces CV death and HF progression when added to the standard therapy in HF patients with reduced ejection fraction, regardless of the presence or absence of diabetes. As such, these trial findings may likely expand the clinical use of SGLT2 inhibitors beyond diabetes care. Nevertheless, the mechanisms underlying the unprecedented benefits of gliflozins in HF management remain elusive.

The presence of renal dysfunction portends adverse outcomes in HF patients. HF is often associated with sodium and water retention, a reduction in renal blood flow and glomerular filtration rate, renal tubular damage, and proteinuria^7, 8^ Despite its cardioprotective actions, SGLT2 inhibitors have also been shown to confer renoprotection by preserving glomerular function, delaying progression to dialysis, reducing urinary protein excretion, and decreasing renal damage in diabetic patients and rodent models^9–11^. However, it remains to be established whether gliflozins are capable of ameliorating renal function in the setting of nondiabetic HF.

The natriuresis elicited by gliflozins, with consequent hemodynamic improvements, constitutes a plausible mechanism underpinning the positive cardiorenal outcomes of these drugs^12^. Interestingly, although SGLT2 inhibitors cause a persistent 7% reduction in the extracellular volume of T2D patients, mice lacking SGLT2 do not exhibit volume depletion, suggesting that the blockade of SGLT2-mediated sodium reabsorption *per se* is not sufficient to affect sodium balance and extracellular fluid homeostasis. In this regard, by coupling a mathematical model of renal function and volume homeostasis with clinical data, Hallow and colleagues predicted that inhibition of apical PT Na^+^/H^+^ exchanger isoform 3 (NHE3) is required for the gliflozin-induced natriuretic effect^13^. In line with the *in silico* predictions, we previously demonstrated that the nonselective SGLT inhibitor phlorizin remarkably reduces PT NHE3 activity^14^. We also found that SGLT2, but not SGLT1, colocalizes with NHE3 in the apical membrane of PT cells^14^. However, the influence of selective SGLT2 inhibitors on NHE3 activity under physiological or pathophysiological conditions has yet to be evaluated.

Based on these observations, the present study aimed to test the hypothesis that an SGLT2 inhibitor is capable of exerting renoprotective effects in the setting of nondiabetic HF. More specifically, we investigated whether empagliflozin improves renal salt and water handling in rats with HF, and we sought to elucidate the potential mechanisms. Furthermore, we examined whether selective SGLT2 inhibition is capable of downregulating PT NHE3 in HF.

## RESULTS

### Empagliflozin lowers serum BNP levels and reduces the right ventricle and lung weight to tibia length ratio in nondiabetic HF rats

The levels of serum BNP higher than 1.0 ng/ml in conjunction with a FAC lower than 40% on echocardiogram were used to characterize HF in rats subjected to myocardial infarction. To this end, blood samples were collected via infraorbital and echocardiographic analysis was undertaken 4 weeks after ligation of the anterior descending coronary artery or sham operation (pretreatment). To determine the effects of empagliflozin on serum BNP and FAC in rats that developed HF, the serum BNP level and echocardiographic analyses were repeated 4 weeks after treatment with empagliflozin or no treatment (posttreatment). As shown in Figure 1A and 1C, the two groups of HF rats had average serum BNP levels of 1.38 ± 0.08 and 1.44 ± 0.12 ng/ml and FAC of 24 ± 2% and 22 ± 2% in the pretreatment period; only rats that achieved the abovementioned criteria were included in the study. At posttreatment, untreated HF rats displayed an increase in serum BNP levels compared with the pretreatment period (2.13 ± 0.25 vs. 1.38 ± 0.08, P < 0.001) (Figure 1A-B). Conversely, HF rats treated with the SGLT2 inhibitor empagliflozin exhibited reduced serum BNP levels compared with the pretreatment period (0.87 ± 0.06 vs. 1.44 ± 0.12, P < 0.001) (Figure 1A-B). Importantly, treatment with empagliflozin restored serum BNP to levels similar to those of sham animals (Figure 1B). A significant decrease in FAC was observed in untreated HF rats from the pretreatment period to the posttreatment period (Figure 1C-D), which in combination with the increase in serum BNP, indicates that cardiac dysfunction deteriorated further in this experimental group. In contrast, empagliflozin treatment induced a modest but significant improvement in FAC during this same time interval (Figure 1C-D). As expected, serum BNP levels and FAC were similar between untreated and empagliflozin-treated sham rats and did not differ between the pretreatment and posttreatment periods.

**Figure 1.**
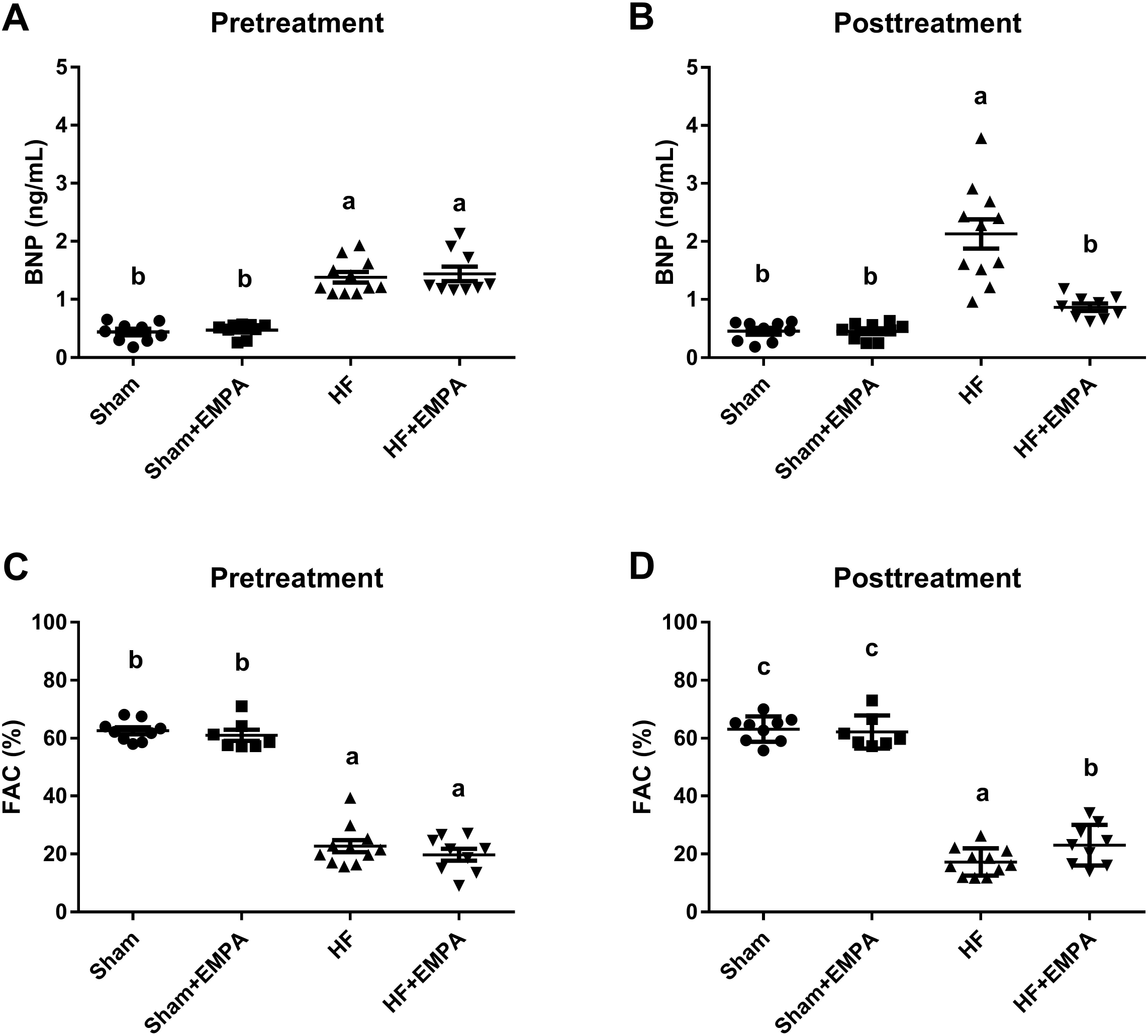
Treatment with empagliflozin (EMPA) normalizes serum brain natriuretic peptide (BNP) concentration and slightly improves LV systolic function in nondiabetic rats with induced heart failure (HF). Doppler echocardiography and quantitative determination of serum BNP were performed 4 weeks after myocardial infarction for HF induction or sham surgery (pretreatment) and after 4 weeks of treatment with EMPA or no treatment (posttreatment). The circulating levels of BNP measured in HF and sham rats at **(A)** pretreatment and **(B)** posttreatment. The fractional area change (FAC) in HF and sham rats at **(C)** pretreatment and **(D)** posttreatment. The values represent individual measurements and the means ± SEM. Different lowercase letters in the scatter plots represent significant differences (P < 0.05).

The biometric characteristics of the animals are shown in Table 1. The average body weight gain was similar among the untreated HF, empagliflozin-treated HF and empagliflozin-treated sham groups (but not the untreated sham group). Therefore, organ weight was normalized by tibial length, which remained unchanged among the four groups of rats. The biometric analysis showed that untreated HF rats exhibited a higher right ventricle (RV) weight to tibia length ratio, a higher lung to tibia length ratio, and more pulmonary congestion than the sham groups. In contrast, empagliflozin prevented pulmonary congestion and RV hypertrophy (Table 1).

**Table 1.**
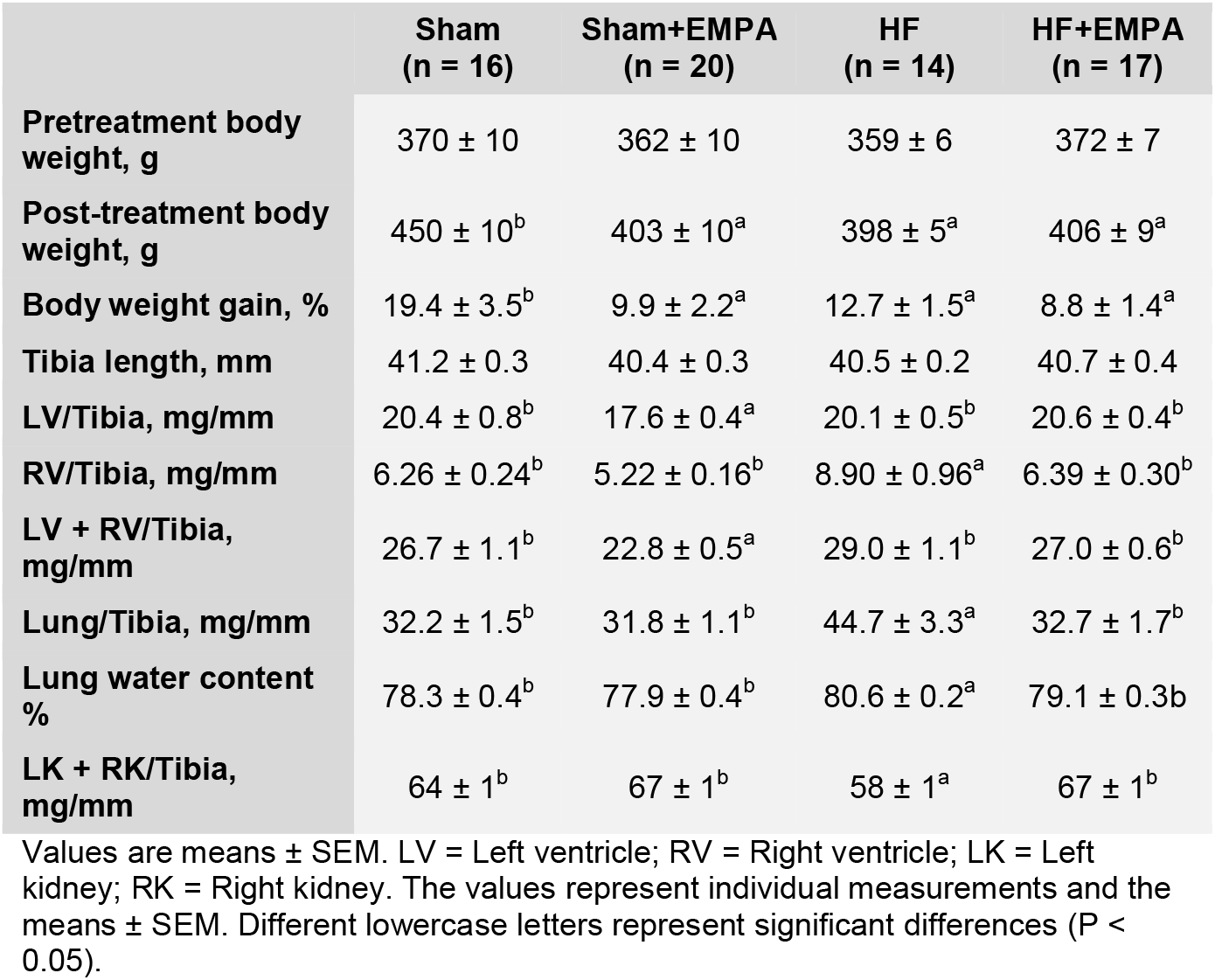
Biometric parameters of sham and non-diabetic rats with myocardial induced-heart failure (HF) treated with the SGLT2 inhibitor empagliflozin (EMPA) or untreated

### Glycosuria, diuresis, and natriuresis induced by empagliflozin were higher in HF rats than in sham rats

We measured urinary excretion of glucose and sodium and urinary flow during the posttreatment period to confirm that continuous administration of empagliflozin (supplied at 10 mg/kg/day in the rat chow) was indeed associated with the suppression of renal glucose, sodium, and water reabsorption. As seen in Figure 2, these parameters were also measured during the pretreatment period. As expected, glycosuria was low and similar among the four groups of rats during the pretreatment period (Figure 2A) but markedly increased in both the sham and HF groups treated with empagliflozin compared with untreated animals (Figure 2B).

**Figure 2.**
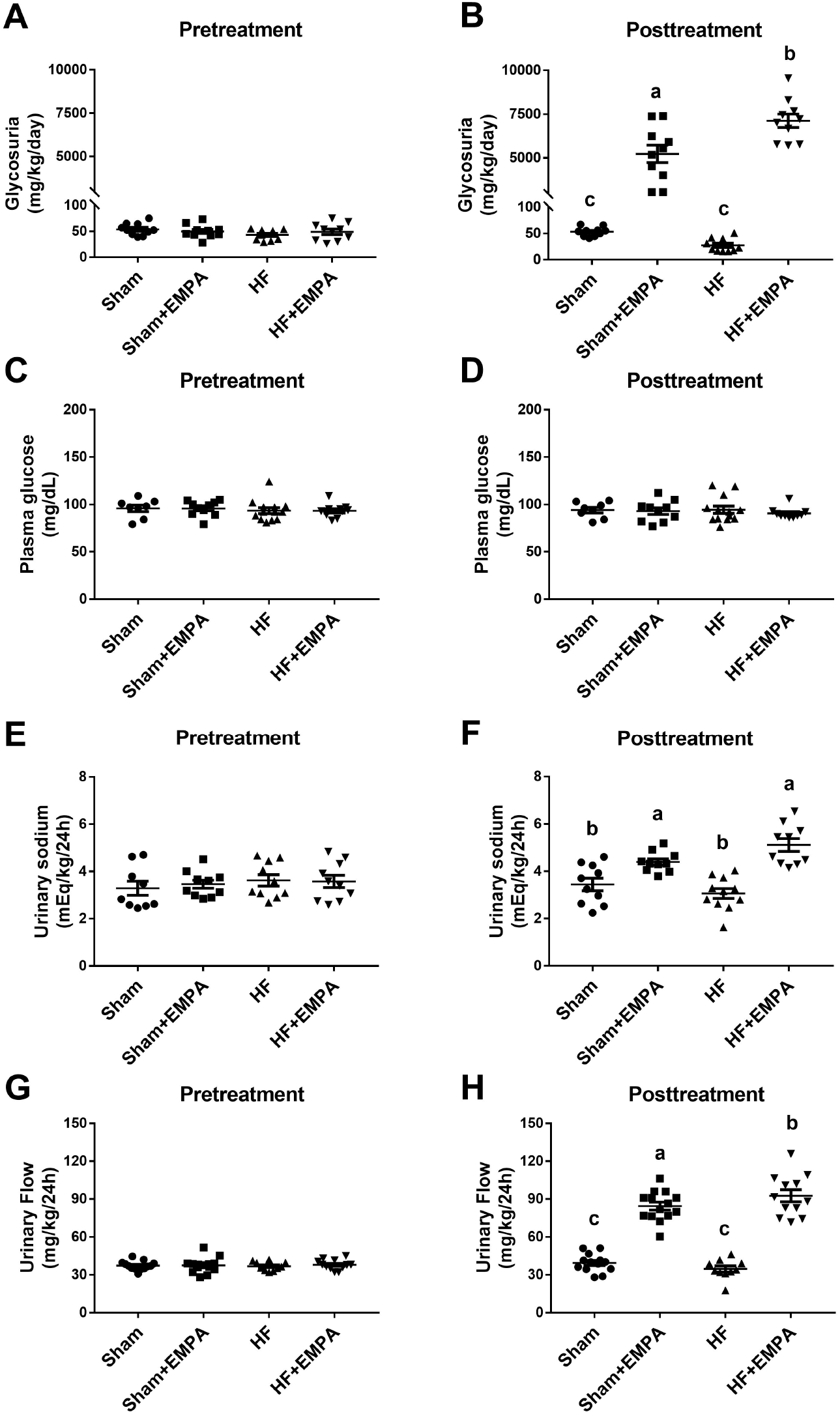
Treatment with empagliflozin (EMPA) induces higher glycosuria, urinary flow, and sodium excretion in HF rats than in sham rats. Rats were individually placed into metabolic cages for 24-h urine collection to measure glycosuria, urinary flow, and urinary sodium. Blood was collected from 12-h fasting rats for the determination of serum glucose. **(A-B)** Glycosuria. **(C-D)** Serum glucose concentration. **(E-F)** Urinary flow. **(G-H)** Urinary sodium excretion. Experiments were conducted 4 weeks after myocardial infarction for HF induction or sham surgery (pretreatment) and 4 weeks after treatment with EMPA or no treatment (posttreatment). The values represent individual measurements and the means ± SEM. Different lowercase letters in the scatter plots represent significant differences (P < 0.05).

Interestingly, the effect of empagliflozin on glycosuria was more pronounced in HF rats than in sham rats (7,118 ± 387 vs. 5,231 ± 499 mg/kg/day, P < 0.01) (Figure 2B). Changes in urinary glucose excretion among the four groups of rats were not associated with changes in serum glucose concentration (Figure 2C-D). Similar to the findings of urinary glucose excretion, no differences were found in urinary flow (Figure 2E) or urinary sodium excretion (Figure 2G) among any of the groups during the pretreatment period. In contrast, during posttreatment, urinary flow (Figure 2F) and sodium excretion (Figure 2H) were higher in the empagliflozin-treated sham group and the HF group than in the untreated groups. Moreover, urinary sodium excretion was greater in empagliflozin-treated HF rats than in empagliflozin-treated sham animals. The increase in urinary flow and urinary sodium excretion was accompanied by increased water and sodium intake (Figure S2).

### SGLT2 is overexpressed in the proximal tubule (PT) of nondiabetic HF rats

The finding that the glycosuric and natriuretic effects of empagliflozin are more pronounced in HF rats than in sham rats suggests that the PT activity of SGLT2 is increased in nondiabetic HF. Therefore, we tested the hypothesis that higher PT SGLT2 activity would be accompanied by increased SGLT2 expression in this nephron segment. SGLT2 protein expression in the PT of the four groups of rats was first evaluated by immunohistochemistry. Representative photomicrographs of SGLT2-stained sections of the renal proximal tubules of HF and sham rats are presented in Figure 3A. The results of this qualitative analysis strongly suggested that SGLT2 was upregulated in nondiabetic HF rats compared with sham rats and HF rats treated with empagliflozin (Figure 3A).

**Figure 3.**
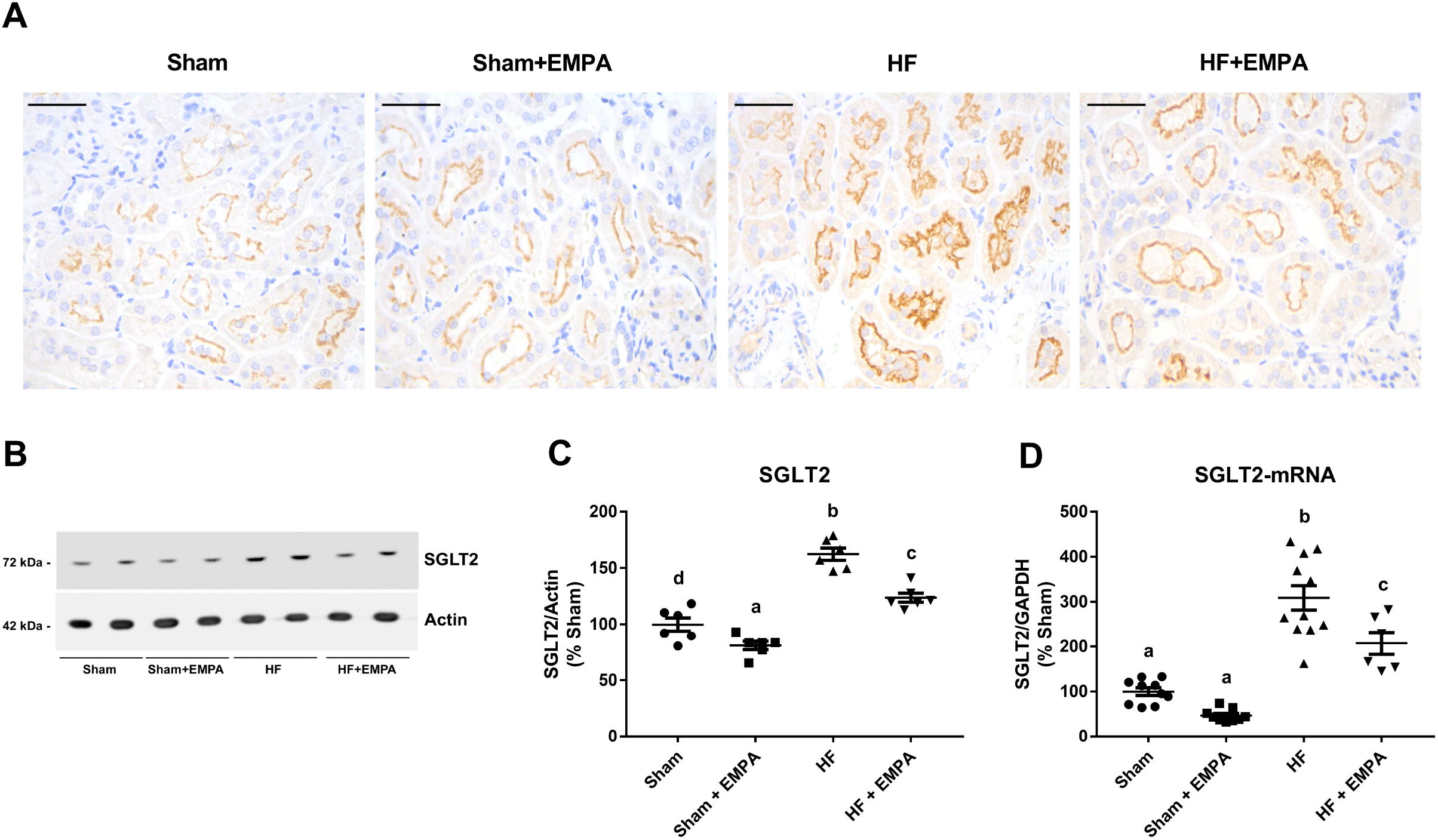
Sodium-glucose cotransporter type 2 (SGLT2) is overexpressed in the proximal tubule (PT) of nondiabetic heart failure (HF) rats. **(A)** Representative immunohistochemical staining of SGLT2 in the PT of sham and HF rats treated with empagliflozin (EMPA) or untreated. Scale bar: 50 μm. **(B)** Representative immunoblots from SDS-PAGE of renal cortical proteins isolated from sham and HF rats treated with EMPA or untreated and probed with antibodies against SGLT2 and actin. **(C)** Graphical representation of the relative levels of SGLT2 protein abundance in the renal cortex of the four groups of rats. **(D)** Graphical representation of the relative mRNA expression of SGLT2 in the renal cortex of sham and HF rats treated with EMPA or untreated. The levels of SGLT2 mRNA were measured using quantitative PCR, and GAPDH was used as an internal control. The values represent individual measurements and the means ± SEM. Different lowercase letters in the scatter plots represent significant differences (P < 0.05).

Second, SGLT2 protein abundance was evaluated in the renal cortex by immunoblotting. As seen in Figure 3B-C, SGLT2 protein abundance was enhanced in the renal cortex of HF rats compared with both untreated and empagliflozin-treated sham rats.

Finally, we found that higher SGLT2 protein abundance was accompanied by higher levels of SGLT2 mRNA expression in the renal cortices of HF rats compared with sham animals (Figure 3D). Interestingly, empagliflozin treatment lowered SGLT2 expression at both the protein and mRNA levels compared with untreated HF rats (Figure 3B-D). However, SGLT2 expression remained higher in empagliflozin-HF rats than in sham rats (Figure 3B-D).

### Empagliflozin restores extracellular volume homeostasis in nondiabetic HF rats

First, we tested the acute renal capability of handling salt and water of HF and sham rats that were either treated with empagliflozin or untreated by challenging these animals with an intraperitoneal bolus of warm saline (10% volume/body weight) and immediately placing them in metabolic cages for 3 hours to collect urine (Figure 4). The percentage of the fluid and sodium load that was excreted within 3 hours of the saline challenge was similar among the four groups of rats during the pretreatment period (Figure 4A and 4C). However, posttreatment, untreated HF rats excreted less fluid (Figure 4B) and salt (Figure 4D) than sham rats and empagliflozin-treated HF rats. Similar fluid and salt load percentages were excreted by sham- and empagliflozin-treated HF rats (Figure 4B and D). In contrast, the acute diuretic and natriuretic responses of empagliflozin-treated sham rats were significantly higher than those of all other groups (Figure 4B).

**Figure 4.**
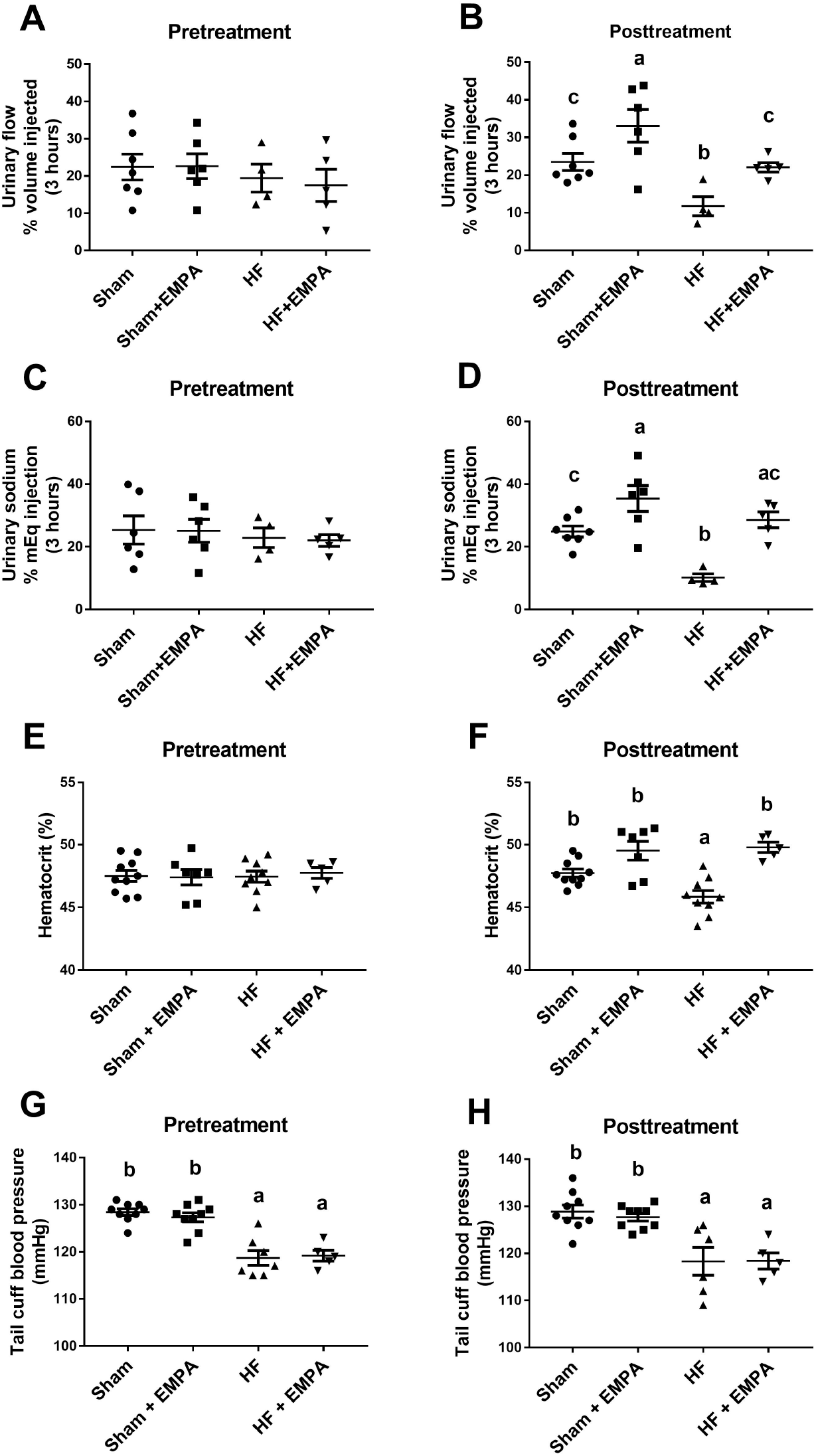
Treatment with empagliflozin (EMPA) improves the volume homeostasis in nondiabetic heart failure (HF) rats. **(A-D)** Rats were challenged with an intraperitoneal bolus of warm saline equivalent to 10% of their body weight and were then placed in metabolic cages for 3-h urine collection. **(A-B)** The percentage of the fluid load that was excreted within 3 hours of the saline challenge. **(C-D)** The percentage of the sodium load that was excreted within 3 hours of the saline challenge. **(E-F)** Blood samples were collected to measure hematocrit. **(G-H)** Blood pressure was measured using tail-cuff plethysmography. Experiments were conducted four weeks after myocardial infarction for HF induction or sham surgery (pretreatment) and four weeks after treatment with EMPA or no treatment (posttreatment). The values represent individual measurements and the means ± SEM. Different lowercase letters in the scatter plots represent significant differences (P < 0.05).

Second, changes in hematocrit were evaluated as a surrogate for changes in volume status. No difference in hematocrit levels was observed among the four groups of rats at pretreatment (Figure 4E). On the other hand, posttreatment, the hematocrit levels of HF rats were lower than those of sham rats (Figure 4F). In addition, treatment with empagliflozin restored the hematocrit of HF rats to levels similar to those of sham rats (Figure 4F).

Third, as seen in Figure 4G-H, the effects of empagliflozin on reducing extracellular volume in HF rats were not accompanied by reduced blood pressure. Untreated and empagliflozin-treated HF rats exhibited lower tail-cuff blood pressure than sham rats during both the pretreatment and posttreatment periods (Figure 4G-H). In addition, empagliflozin treatment did not affect blood pressure in either sham or HF rats (Figure 4H).

### Empagliflozin prevents the reduction in GFR, attenuates proteinuria, and preserves kidney mass in nondiabetic HF rats

As depicted in Figure 5, renal function parameters were evaluated during both the pretreatment and posttreatment periods. To this end, rats were individually placed in metabolic cages for 24-h urine collection. In the pretreatment period, no differences were observed in GFR (Figure 5A) or urinary protein excretion among the four groups of rats (Figure 5E), except for higher plasma urea levels in HF rats than in sham rats (Figure 5C). Nevertheless, posttreatment, HF rats exhibited lower GFR (Figure 5B), higher levels of plasma urea (Figure 5D), and greater proteinuria (Figure 5F) than the other three groups of animals. In addition, untreated HF rats displayed a deterioration in these renal function parameters from the pretreatment to the posttreatment period (Figure 5A-F). In contrast, empagliflozin-treated HF rats exhibited GFR, serum urea, and proteinuria similar to sham groups, suggesting that empagliflozin prevented and/or restored renal function in HF rats. Interestingly, HF rats exhibited a lower kidney weight to tibia length ratio than sham rats, whereas empagliflozin treatment prevented HF-induced kidney atrophy (Table 1).

**Figure 5.**
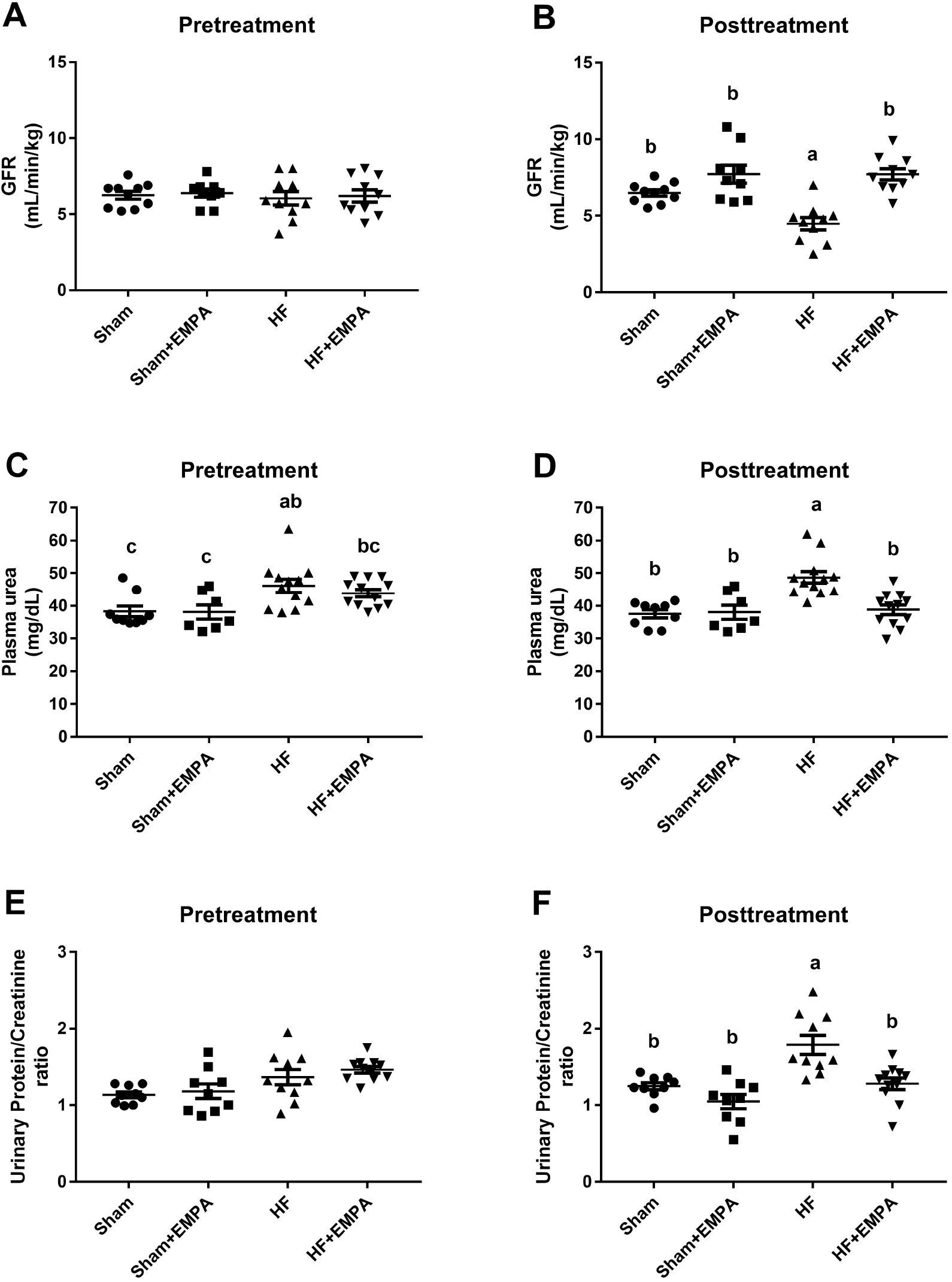
Treatment with empagliflozin (EMPA) prevents the reduction in glomerular filtration rate (GFR) and attenuates proteinuria in heart failure (HF) rats. Rats were individually placed into metabolic cages for 24-h urine collection to measure GFR and proteinuria. Blood was collected for the determination of serum urea and creatinine. **(A-B)** The GFR was estimated by creatinine clearance. **(C-D)** Plasma urea was measured by colorimetry. **(E-F)** Graphical representation of the urine protein to creatinine ratio. Proteinuria was measured by colorimetry. Experiments were conducted 4 weeks after myocardial infarction for HF induction or sham surgery (pretreatment) and 4 weeks after treatment with EMPA or no treatment (posttreatment). The values represent individual measurements and the means ± SEM. Different lowercase letters in the scatter plots represent significant differences (P < 0.05).

### Empagliflozin inhibits NHE3 activity in the renal proximal tubule

We employed stationary in vivo microperfusion to determine NHE3 activity in the proximal tubule to test the hypothesis that selective SGLT2 inhibition is capable of downregulating PT NHE3 in HF. As illustrated in Figure 6A, PT NHE3 activity, evaluated as the rate of bicarbonate reabsorption (J_HCO3-_), was higher in untreated HF rats than in untreated sham rats. Treatment with empagliflozin prominently reduced NHE3 activity in the PT of both HF and sham rats (Figure 6A). Nevertheless, the inhibitory effect of empagliflozin on NHE3 function was more pronounced in HF rats (~ 65% inhibition vs. untreated HF rats) than in sham rats (~ 40% inhibition vs. untreated sham rats). In addition, no differences were observed in NHE3 activity among the four groups of rats when their PTs were perfused with the specific NHE3 inhibitor S3226, confirming that differences in net bicarbonate reabsorption induced by empagliflozin were due to NHE3 (Figure 6B). The increase in PT NHE3 activity in HF rats was accompanied by enhanced NHE3 mRNA levels (Figure 6C) and NHE3 protein abundance (Figure 6D-E). Conversely, even though empagliflozin reduced PT NHE3 activity, this SGLT2 inhibitor did not affect either cortical NHE3 protein abundance or the mRNA levels of NHE3 in HF or sham rats (Figure 6C-E). NHE3 phosphorylation at serine 552 (PS552-NHE3) is considered a surrogate for NHE3 inhibition^16–18^. Consistent with higher PT NHE3 activity, HF rats displayed lower levels of PS552-NHE3 to total NHE3 than the other three groups of rats (Figure 6D and 6F). The levels of PS552-NHE3 to total NHE3 were higher in empagliflozin-treated HF rats than in untreated HF rats but lower than in the rats in the sham groups. Importantly, although empagliflozin inhibited NHE3 activity in the PT of sham rats, no differences were found in the levels of PS552-NHE3 to total NHE3 between untreated and empagliflozin-treated sham rats (Figure 6D and 6F).

**Figure 6.**
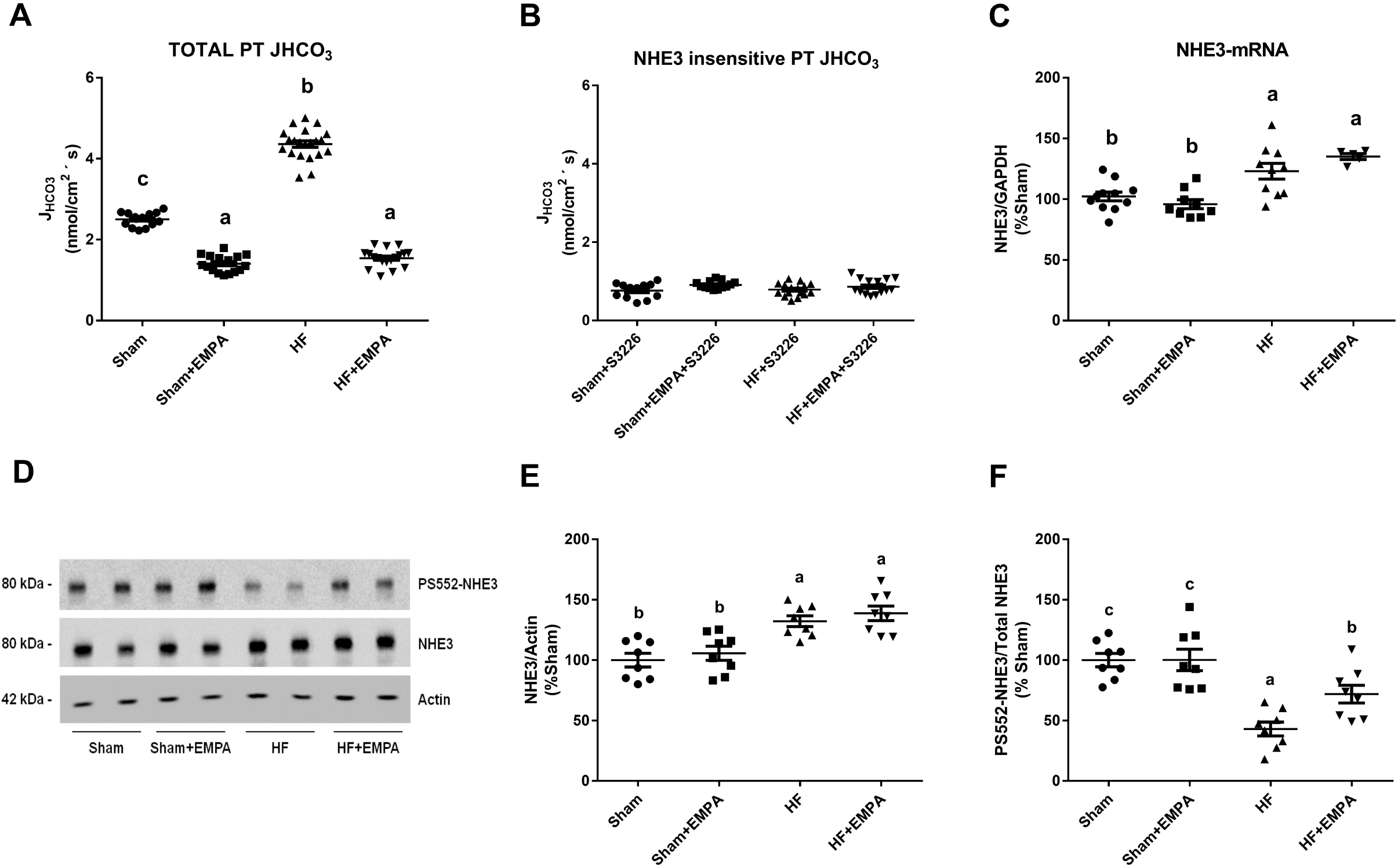
Treatment with empagliflozin (EMPA) inhibits proximal tubule (PT) NHE3 activity. **(A-B)** NHE3 activity was determined as the rate of PT bicarbonate reabsorption. PT bicarbonate reabsorption (J_HCO3-_) in HF and sham rats treated with EMPA or untreated was assessed by stationary microperfusion and continuous measurement of luminal pH in the absence **(A)** or presence **(B)** of 10 μM S3226, a selective inhibitor of NHE3. **(C)** Graphical representation of the relative mRNA expression of NHE3 in the renal cortex of sham and HF rats treated with EMPA or untreated. The levels of NHE3 mRNA were measured using quantitative PCR, and GAPDH was used as an internal control. **(D)** Renal cortical proteins isolated from the four groups of rats were resolved by SDS-PAGE and transferred to PVDF membranes. The membranes were probed with a primary antibody against total NHE3, a phosphospecific antibody that recognizes NHE3 only when it is phosphorylated at serine 552 (PS552-NHE3), and an antibody against actin as an internal control. **(E)** Graphical representation of the relative expression of total NHE3 and **(F)** the ratio of phosphorylated NHE3 at serine 552 to total NHE3 (PS552-NHE3/total). The values represent individual measurements and the means ± SEM. Different lowercase letters in the scatter plots represent significant differences (P < 0.05).

## DISCUSSION

SGLT2 inhibitors reduce the risk of severe HF events, even though SGLT2 is not expressed in the human heart. The present study provides novel experimental evidence that SGLT2 inhibition by empagliflozin exerts renal benefits that may contribute to HF progression in a nondiabetic setting. Our results demonstrate that the mechanisms underlying empagliflozin benefit include the preservation of glomerular function and a profound reduction of PT NHE3-mediated sodium reabsorption in an experimental model of myocardial infarction-induced HF. These effects were accompanied by restoration of extracellular volume homeostasis, a decrease in circulating levels of BNP, prevention of pulmonary congestion, and mitigation of RV hypertrophy. Moreover, to the best of our knowledge, this is the first report to reveal that SGLT2 is overactive and overexpressed in the renal PT of nondiabetic HF rats. Interestingly, empagliflozin not only suppressed SGLT2-mediated glucose reabsorption but also reduced SGLT2 expression in the renal cortex of nondiabetic HF rats at both the protein and mRNA levels.

Excessive renal sodium avidity and extracellular volume overload are hallmark features of HF that are associated with disease progression and poor prognosis^19^. The sodium and water retention that occurs with advanced LV failure can lead to pulmonary edema, increased mean pulmonary arterial pressure, and consequently, RV hypertrophy that is ultimately followed by dysfunction^20^. Accordingly, we found that untreated HF rats exhibited a lower natriuretic and diuretic response to saline challenge, a lower hematocrit (which may indicate lower hemoconcentration as a consequence of intravascular volume expansion) and much higher circulating levels of BNP than sham rats. As expected, volume overload in untreated HF rats was accompanied by pulmonary congestion and RV hypertrophy. Conversely, treatment with empagliflozin restored extracellular volume homeostasis in HF rats, as evidenced by the improved sodium and water handling by the kidneys and normalization of hematocrit and serum BNP levels. The reduction in volume overload by empagliflozin prevented pulmonary congestion and RV hypertrophy in nondiabetic HF rats. In line with these findings, Chowdhury and colleagues^21^ found that empagliflozin treatment in rats with severe experimental pulmonary hypertension is capable of reducing mean pulmonary artery pressure, RV systolic pressure, and RV hypertrophy. Importantly, these hemodynamic benefits were associated with prolonged survival in this model of pulmonary arterial hypertension. These data support the hypothesis that reduced volume overload is one of the mechanisms underlying the observed decrease in HF events in cardiovascular trials with SGLT2 inhibitors.

The potential causes of impaired sodium and water handling by the kidneys in nondiabetic HF may include a decreased GFR, increased tubular reabsorption of sodium, or both. This study demonstrates that empagliflozin prevents a decrease in GFR, diminishes serum urea levels, and exerts antiproteinuric effects in nondiabetic experimental HF. Interestingly, we report the unprecedented finding that treatment with empagliflozin prevents HF-induced renal atrophy. Reduced renal blood flow due to decreased cardiac output and increased venous congestion is probably the primary mediator of GFR decline and loss of renal mass in HF. However, it remains to be determined whether SGLT2 inhibitors are capable of improving renal perfusion in the setting of nondiabetic cardiac dysfunction. Interestingly, treatment with the SGLT2 inhibitor luseogliflozin prevented endothelial rarefaction, renal hypoxia, and the development of renal fibrosis after renal ischemia/reperfusion in mice through a vascular endothelial growth factor (VEGF)-dependent pathway^22^. Furthermore, using a systemic molecular approach, Pirklbauer and colleagues^23^ provided evidence that SGLT2 inhibition blocks the expression of key mediators of renal fibrosis and kidney disease progression in two distinct lines of human proximal tubule cells. It could therefore be speculated that the renoprotective effects of SGLT2 in nondiabetic HF are mediated not only by improvements in hemodynamics but also by local actions at the level of PT cells.

The upregulation of NHE3 expression and renal PT activity have been implicated in the pathogenesis of HF, which includes sodium retention, volume overload, and peripheral edema^24–26^. Consistent with previous findings from our group^27^, we observed increased NHE3 activity, protein abundance, and mRNA levels and decreased PS552-NHE3 in the PT of HF rats. Importantly, in this report, we show for the first time that a selective SGLT2 inhibitor suppressed the activity of PT NHE3. Empagliflozin treatment prominently reduced PT NHE3 function in both HF and sham rats; however, NHE3 inhibition was more pronounced in HF rats. Empagliflozin-induced NHE3 inhibition did not affect NHE3 protein abundance or NHE3 mRNA levels in either sham or HF rats. In contrast, a small but significant increase in PS552-NHE3 levels was observed in empagliflozin-treated HF rats compared with untreated HF rats. As no difference was found in the levels of PS552-NHE3 between untreated and empagliflozin-treated sham rats, one may speculate that the effect of empagliflozin on PS552-NHE3 in HF rats might be secondary to the attenuation of the maladaptive neurohormonal response in HF. Indeed, both activation of the renal sympathetic nervous system and angiotensin II (Ang II) are known to activate NHE3, at least in part, due to a decrease in PS552-NHE3^28, 29^ Nevertheless, the principal mechanism by which SGLT2 inhibition downregulates PT NHE3 activity remains unknown.

The PT reabsorbs two-thirds of the salt and water filtered by the glomeruli, and NHE3 is the primary transporter driving transcellular sodium reabsorption at this nephron segment^30^. As such, NHE3 plays an essential role in extracellular volume homeostasis and blood pressure control. Indeed, PT-specific deletion of NHE3 lowers blood pressure in mice under basal conditions and markedly reduces the increase in blood pressure in response to chronic injections of Ang II compared with wild-type mice^31^. Notwithstanding, the current study demonstrates that empagliflozin-induced inhibition of PT NHE3 in HF rats was followed by reduced volume overload but not by decreased blood pressure. A potential explanation for these findings is that SGLT2 inhibition may indirectly upregulate sodium-mediated reabsorption in the distal nephron, partially compensating for the diminished sodium reabsorption by the PT. In line with this hypothesis, the SGLT2 inhibitor luseogliflozin significantly increases the phosphorylated levels of the sodium chloride cotransporter (NCC), a surrogate for active NCC, in the renal cortex of obese rats^32^.

SGLT2 expression and activity appear to be upregulated in PT cells of patients with T2D as a result of a maladaptive regulatory mechanism that contributes to the maintenance of hyperglycemia^33^, Here, we showed that the glycosuric effects of empagliflozin were higher in empagliflozin-treated HF rats than in empagliflozin-treated sham rats, indicating that SGLT2 activity is upregulated in the PT of nondiabetic HF animals. Overactive SGLT2 in HF is due to increased levels of SGLT2 protein abundance and SGLT2 mRNA levels. The upregulation of SGLT2 expression in nondiabetic HF may involve hyperactivation of the sympathetic nervous system and/or the renal angiotensin system (RAS). In this regard, Matthews and colleagues revealed that supraphysiological concentrations of norepinephrine increase the total abundance and surface expression of SGLT2 in the apical membrane of human PT cells exposed to normal glucose levels^35^. SGLT2 upregulation by Ang II has also been observed in nondiabetic settings, such as in renovascular hypertension^36^. Sodium-dependent glucose uptake, SGLT2 protein abundance, and transcription were increased in the PT of renovascular hypertensive rats. In addition, when these secondary hypertensive rats were treated with ramipril or losartan, the increases in both blood pressure and activity/expression of SGLT2 were mitigated^36^. Notably, in vitro studies have suggested that the relationship between SGLT2 and RAS may be bidirectional, since increased glucose uptake by PT leads to RAS activation^37, 38^, The vicious cycle of SGLT2 upregulation and intrarenal RAS hyperactivation may aggravate disorders of extracellular fluid volume and therefore contribute to HF progression. Likewise, gliflozins may break this vicious cycle by lowering PT glucose uptake and decreasing RAS activation, thereby mitigating the potential stimuli for SGLT2 overexpression.

In summary, our findings demonstrated that empagliflozin confers therapeutic benefits in nondiabetic HF rats by restoring the euvolemic status, most likely due to the preservation of GFR and the inhibition of NHE3-mediated sodium reabsorption. Furthermore, our study provides novel evidence that SGLT2 overactivity and overexpression may constitute a mechanism for PT dysfunction implicated in the development and progression of HF syndrome. Aligned with recent clinical data showing that dapagliflozin confers cardioprotection in nondiabetic HF patients^6^, our report supports the recommendation of SGLT2 inhibitors in future treatment paradigms for HF management, independent of diabetes status.

## MATERIAL AND METHODS

A detailed description of the materials and methods can be found in the online supplement.

### Animal protocols, surgical procedures, and drug treatment

All experiments were carried out following the ethical principles in animal research of the Brazilian College of Animal Experimentation and were approved by the Institutional Animal Care and Use Committee (Protocol # 941/2018). Myocardial infarction (MI) was induced in 8-week-old male Wistar rats (200-250 g) (n = 50) by ligation of the left anterior descending (LAD) artery, as previously described^15^. Age-matched sham-operated animals underwent an identical surgical procedure without actual LAD artery ligation (n = 44). Four weeks after surgery (pretreatment), MI rats that developed HF and sham rats were randomly divided into two groups and treated for four weeks with empagliflozin (10 mg/kg/day supplied in the rat chow, posttreatment). At pretreatment, HF was characterized by echocardiographic evaluation of left ventricle (LV) systolic function and serum levels of brain natriuretic peptide (BNP). HF was considered when the fractional area change (FAC) was lower than 40% and when circulating levels of BNP were higher than 1.0 ng/ml. Blood pressure measurements, renal function evaluation, and saline challenge were performed during both the pretreatment and posttreatment periods. The rats were anesthetized by an intraperitoneal injection of pentobarbital (50 mg/kg) and subsequently killed by decapitation. Arterial blood was collected at the time of death. The kidneys, heart, and lungs were excised and weighed. The kidneys were immediately removed for isolation of renal cortical proteins and immunoblotting, for tissue fixation for immunohistochemistry, or for freezing for RNA extraction and quantitative RT-PCR. Stationary in vivo microperfusion was employed to measure PT NHE3 activity in 4-5 rats/group. The study design is depicted in Supplementary Figure S1.

### Statistical analysis

The results are reported as the mean ± standard error of the mean (SEM), with *n* indicating the number of rats. Comparisons among the means were assessed using a two-way analysis of variance (ANOVA) followed by the Newman-Keuls post hoc test. A *P* value < 0.05 was considered significant, taking into account the main effect of empagliflozin treatment, the main effect of heart failure induction, the interaction between empagliflozin treatment and heart failure induction, and the differences among groups.

## SOURCES OF FUNDING

This work was supported by the São Paulo State Research Foundation (FAPESP) Grant 2016/22140-7, by the Brazilian National Council for Scientific and Technological Research (CNPq), and by the Coordination for the Improvement of Higher Education Personnel (CAPES).

## Nonstandard Abbreviations and Acronyms

Ang II: angiotensin II
BNP: brain natriuretic peptide
NCC: sodium chloride cotransporter
CV: cardiovascular
DAPA-HF: **D**apagliflozin **A**nd **P**revention of **A**dverse-outcomes in **H**eart **F**ailure
EMPA: empagliflozin
FAC: fractional area of change
GFR: glomerular filtration rate
HF: heart failure
J_HCO3-_: rate of bicarbonate reabsorption
LAD: left anterior descending
LV: left ventricle
MI: myocardial infarction
NHE3: Na^+^/H^+^ exchanger isoform 3
PS552-NHE3: NHE3 phosphorylated of at the serine 552
PT: proximal tubule
RAS: renal angiotensin system
RV: right ventricle
SGLT1: sodium-glucose cotransporter type 1
SGLT2: sodium-glucose cotransporter type 2
T2D: type 2 diabetes
VEGF: vascular endothelial growth factor

## DISCLOSURES

The authors have declared no conflicts of interest.

## Notes

### Competing Interest Statement

The authors have declared no competing interest.

