## Supplemental for "Empagliflozin inhibits proximal tubule NHE3 activity, preserves GFR and restores euvolemia in nondiabetic rats with induced heart failure"

### SUPPLEMENTAL METHODS

*Reagents and antibodies* - Jardiance tablets (Boehringer Ingelheim Pharma GmbH & Co. KG) containing 25 mg of empagliflozin were purchased from a local pharmacy. These tablets were then sent to the Rhoster Company (Araçoiaba da Serra, São Paulo, Brazil) to be added to the rodent chow for daily empagliflozin treatment (10 mg/kg/day). Untreated animals received a control diet without empagliflozin. The monoclonal antibody against NHE3[1] (1:1,000 dilution) was a gift from Peter Aronson, Yale University School of Medicine (New Haven, CT, USA). The monoclonal antibody against PS552-NHE3[1] (1:1,000 dilution) was purchased from Santa Cruz Biotechnology (sc-53962, Santa Cruz, CA, USA). The polyclonal antibody against SGLT2 (1:1,000 and 1:100 dilution for immunoblotting and immunohistochemistry, respectively) was purchased from Novus Biologicals (NBP1-92384, Centennial, CO, USA). The monoclonal antibody against actin (1:5,000 dilution) was acquired from Abcam (ab179467, Cambridge, MA, USA). Secondary antibodies were from Jackson ImmunoResearch Laboratories, Inc (West Grove, PA, USA). Chemicals were obtained from Merck (Darmstadt, Germany) unless otherwise specified.

*Animals protocols, surgical procedures, and drug treatment* - Eight-week-old male Wistar rats (200-250g) (n = 104) were obtained from the University of São Paulo Medical School, São Paulo, SP, Brazil. Rats were randomly assigned to sham surgery (n = 36) or myocardial infarction (MI) (n = 68) by ligation of the left anterior descending (LAD) artery (Supplemental Figure S1), as previously described[2]. Briefly, rats were anesthetized with ketamine (50 mg/kg) and xylazine (10 mg/kg) intraperitoneally and placed under positive pressure ventilation (rate: 90 breaths/minute; tidal volume: 2.5 ml on a Harvard rodent respirator [model 683, Harvard Apparatus Co., South Natick, MA, USA]). The thoracotomy was performed at the fourth intercostal space. After that, the heart was exposed, and the LAD coronary artery was isolated and ligated ~3 mm from the origin of the aorta using 6-0 Prolene suture. The chest was closed, and the animals were carefully moved from ventilation support. Sham-operated rats were submitted to a similar surgical procedure, except for LAD coronary artery ligation. After recovering from surgery, the rats were maintained in a temperature and humidity-controlled environment with a 12-hour dark/light cycle at the Heart Institute (InCor) animal facility. Food and water were supplied ad libitum.

Four weeks after surgery (pretreatment), HF was characterized by echocardiographic evaluation of left ventricle (LV) systolic function and serum levels of brain natriuretic peptide (BNP). HF was considered when the fractional area change (FAC) was lower than 40%, and when circulating levels of BNP were higher than 1.0 ng/ml. MI rats that developed HF (n = 39) and sham (n = 36) rats were randomly divided into two groups and treated with empagliflozin [10 mg/kg/day supplied in the rat chow] or no treated (Supplemental Figures S1 and S2). After four weeks (posttreatment), the rats were anesthetized by an intraperitoneal injection of pentobarbital (50 mg/kg) and subsequently killed by decapitation.

*Echocardiography* - Rats were anesthetized with 1.5% isoflurane in $O_2$ and placed in the left lateral decubitus position to obtain cardiac images. Images were captured and analyzed using Sonos 5500 ultrasound equipment (Philips Medical System, Bothell, WA) with a 12-14 MHz transducer (2 cm depth with fundamental and harmonic imaging). Echocardiographic images were acquired by placing the cursor of pulsed-wave Doppler in the LV outflow tract to display the end of aortic ejection and the onset of mitral inflow. Tracing the endocardial border by planimetry, excluding the papillary muscle, in end-diastolic and end-systolic frames provide values for LV end-diastolic area (LVEDA) and LV end-systolic area (LVEDS). Applying this in the following equation would give a FAC of LV from diastole to systole: FAC = [LVEDA – (LVEDS/LVEDA)] x 100%. Echocardiography was performed by an investigator who was blind to the experimental groups.

*Determination of BNP serum levels* - At pretreatment, blood samples were withdrawn from the retro-orbital sinus under isoflurane anesthesia. At posttreatment, blood was collected from the abdominal aorta artery at the time of death. The samples were immediately transferred into vacuum tubes with gel separator (BD vacutainer® SST® II Advance®) and centrifugated at 4,000 rpm for 10 minutes at 4°C to obtain serum. Serum levels of BNP were measured by enzyme-linked immunoassay (ELISA) (BNP 32 Rat ELISA kit, Abcam) according to the manufacturer's instructions.

*Blood pressure measurements* - Tail-cuff blood pressure was measured noninvasively in conscious restrained rats by plethysmography (BP Blood Pressure Analysis System - 2000, Visitech System®, Apex, NC, USA). Before measurements, the rats were acclimated to restraint and tail-cuff inflation for 30 minutes per day for three days.

*Renal function evaluation* - The rats were individually housed and placed in metabolic cages (Tecniplast, Buguggiate, VA, Italy) as previously described[3]. Food and water consumption were determined daily and subsequently normalized to body weight. Urine samples were collected for 24 hours and used to determine urinary flow, sodium, creatinine, and total proteinuria. Creatinine clearance was used to estimate the glomerular filtration rate (GFR).

*Blood and urine analysis* - Fasting plasma glucose levels were measured using the ACCU-CHECK® Performa meter (Roche Diagnostics GmbH, Mannheim, Germany). The serum and urinary sodium concentrations were determined by flame photometry (Digimed® DM-62, São Paulo, Brazil). The plasma urea, urinary glucose, and creatinine concentrations were measured by colorimetric methods using Labtest kits (Labtest, Minas Gerais, Brazil). The urinary protein excretion was determined using a Sensiprot kit (Labtest). The experiments were carried out following the manufacturer's instructions.

*Saline challenge* - Animals were anesthetized with isoflurane and were injected intraperitoneally with a volume of warmed (37°C) saline (0.9% NaCl) equivalent to 10% of their body weight (v/w). After that, the rats were placed immediately in metabolic cages for 3 hours of urine collection. Urine volume was measured with a graduate pipette, and urinary sodium concentration was measured by flame photometry (Digimed® DM-62). The results were expressed as the percentage of the fluid and sodium load that was injected.

*Biometric and morphometric analysis* - The lungs, heart, and kidneys were excised and weighed. The organ weight was normalized by left tibia length. The lungs were stored in an oven to dry at 70°C for 48 hours. The relative water content of lung tissue was calculated using the following equation: Lung water content (in %) = (wet lung weight − dry lung weight)/lung wet weight × 100. The kidneys were immediately removed for isolation of renal cortical proteins for immunoblotting, for tissue fixation for immunohistochemistry, or for freezing for RNA extraction and quantitative RT-PCR.

*Preparation of renal cortical homogenate* - The right kidney from rats was removed, cut in half and the cortices were isolated and homogenized in ice-cold PBS buffer (150 mM sodium chloride, 2.8 mM monobasic sodium phosphate, 7.2 mM dibasic sodium phosphate, pH 7.4) containing protease inhibitors (0.7 µg/ml pepstatin, 0.5 µg/ml leupeptin and 40 µg/ml phenylmethanesulfonyl fluoride) and phosphatase inhibitors (50 mM sodium pyrophosphate decahydrate and 15 mM sodium fluoride) using a Potter-Elvehjem-style tissue grinder (POLIMIX® PX-SR50E, Kinematica Inc., Luzern, Switzerland). The homogenate was centrifuged at 4,500 rpm for 10 minutes at 4°C. The pellets were resuspended in fresh PBS containing protease and phosphatase inhibitors and stored at -80°C. Protein concentration was determined by the Lowry method[4].

*SDS-PAGE and immunoblotting* - Renal cortical proteins were solubilized in Laemmli sample buffer and separated by SDS-PAGE using 7.5% or 10% polyacrylamide gels. For immunoblotting, proteins were transferred to PVDF membranes (Millipore Immobilon-P, Millipore®, Bedford, MA) at 350 mA for 8-10 hours at 4°C with a TE 62 transfer electrophoresis unit (GE HealthCare®). Sheets of PVDF containing transferred proteins were incubated first in Blotto (5% nonfat dry milk and 0.1% Tween 20 in PBS, pH 7.4) for 1 hour to block nonspecific binding, followed by overnight incubation with primary antibodies diluted in Blotto. The sheets were then washed five times in Blotto and incubated for 1 hour at room temperature with a respective appropriated horseradish-peroxidase-conjugated immunoglobulin secondary antibody diluted 1:2,000 in Blotto. After washing five times in Blotto and twice in PBS (pH 7.4), the PVDF membrane was incubated for 1 minute with an enhanced chemiluminescence detection (ECL) system (GE HealthCare®) for visualization of the bound antibodies. The visualized bands were digitized using the ImageScanner III (GE HealthCare®) and quantified using ImageJ software (National Institutes of Health, Bethesda, MD).

*Immunohistochemistry* - The left kidney from rats were cut in half on a midsagittal plane, fixed in 10% formalin for 24 hours, stored in 70% ethanol, and then embedded in paraffin. Four-micrometer kidney paraffin sections were incubated with 3% $H_2O_2$ for 3 minutes (five times at room temperature) to block endogenous peroxidase activity and then rinsed with TBST. Nonspecific reactions were blocked in 2% goat serum for 20 minutes and then incubated with the rabbit polyclonal anti-SGLT2 antibody (1:100 dilution). After 18 hours of incubation at 4°C, kidney sections were washed 3 times for 5 minutes with TBST and incubated with a secondary antibody. After washing in TBST, tissue sections were incubated with peroxidase-conjugated universal immuno-enzyme polymer, anti-rabbit solution (Histofine® Simple Stain™ MAX PO(MULTI), Nichirei Biosciences Inc, Tokyo) for 30 minutes at room temperature. After washing in TBST, immunoreactions were detected with 3,3'-diaminobenzidine tetrahydrochloride (DAB-Zymed®) for 7 minutes and counterstained with hematoxylin. The images were acquired under a 400x magnification light microscope using the software (Quantimet Leica, Leica Biosystems®, Wetzlar, HE).

*Quantitative Real-Time RT-PCR* - Total RNA was isolated from the left kidney using Trizol (Thermo Fisher Scientific, Carlsbad, CA) according to the manufacturer's instructions, quantified (ND-1000 spectrophotometer-NanoDrop Technologies, Inc.), and treated with DNase-I. First-strand cDNA synthesis was performed using Super-Script III Reverse Transcriptase (Invitrogen) following the manufacturer's guidelines. Details about the oligonucleotide primers used in this study are listed in Table S1. PCR products were visualized on 0.8% agarose gels with ethidium bromide. Reactions were carried using SYBR Green PCR Master Mix-PE (Thermo Fisher Scientific) on an ABI Prism® 7500 Fast Sequence Detection System (Applied Biosystem, Foster City, CA). The comparative threshold cycle method was used for data analyses. All samples were assayed in triplicate. Transcripts for three reference genes (G*apdh, Actb,* and *Ppia)* were determined (Supplemental Table S1). The BestKeeper software[5] was used to identify the best suit reference gene (Cyclophilin A) for data normalization under our experimental conditions. Relative expression was analyzed by the $2^{-\Delta\Delta}$CT method.

*In vivo stationary microperfusion* - Rats were anesthetized by intramuscular administration of tiletamine/zolazepam (50 mg/kg) and xylazine (5 mg/kg). After tracheostomy, the left jugular vein was cannulated for infusion of 3% mannitol in isotonic saline solution, at a rate of 0.05 mL/min. The kidney was isolated using a lumbar approach, immobilized in situ using Ringer-agar in a Lucite cup under a microscope, and adequately illuminated. Proximal tubules were punctured using a double-barrelled micropipette, one barrel being used to inject FDC-green colored Ringer perfusion solution (in mM: 100 NaCl, 5 KCl, 25 $NaHCO_3$, 1 $CaCl_2$, 1.2 $MgSO_4$, and raffinose to reach isotonicity), and the other to inject Sudan-black colored castor oil, the latter used to block the injected fluid column in the lumen. To measure luminal pH, proximal tubules were impaled by a double-barreled asymmetric microelectrode, the larger barrel containing H-ion-sensitive ion-exchange resin silanized with hexamethyldisilazane (Sigma

Fluka, Buchs, Switzerland) and the smaller barrel containing the reference solution (1 M KCl) colored by FDC-green. The rate of tubular acidification was measured by injecting a droplet of the perfusion solution between the oil columns and following the luminal pH changes toward the steady-state level (stationary perfusion). The voltage between the microelectrode barrels, representing luminal $H^+$ activity, was continuously recorded with a microcomputer equipped with an analog-to-digital conversion board (Lynx, São Paulo, Brazil) for data acquisition and processing. Luminal bicarbonate was calculated from luminal pH and arterial blood $P_{CO2}$, and the rate of tubular acidification was expressed as the half time of the exponential reduction of the injected $HCO_3^-$ concentration to its stationary level ($t_{1/2}$). Net $HCO_3^-$ reabsorption ($J_{HCO3-}$) per $cm^2$ of tubule epithelium was calculated by using the following equation:

$$J_{HCO3-} = \frac{\ln 2}{t_{1/2}}[(HCO_3^-)_0 - (HCO_3^-)_s]*\frac{r}{2}$$

where k is the rate constant of luminal bicarbonate reduction, [k = ln2/($t_{1/2}$)], r is the tubule radius, and $(HCO_3^-)_0$ and $(HCO_3^-)_s$ are the $HCO_3^-$ concentrations at the infused and stationary level, respectively. The tubules were perfused with a control solution in the presence or absence of the selective NHE3 inhibitor, S3226 (10 μM)[6].

**SUPPLEMENTAL TABLE**

**Table S1. Sequences of oligonucleotides used in this study.**

| Gene | Primer Sequence 5' to 3' | Size (base pairs) |
|---|---|---|
| *Gapdh* | F - ATGGTGAAGGTCGGTGTG<br>R - GAACTTGCCGTGGGTAGAG | 162 |
| *Actb* | F - CGTTGACATCCGTAAAGACC<br>R - GCCACCAATCCACACAGA | 172 |
| *Ppia* | F - AATGCTGGACCAAACACAAA<br>R - CCTTCTTTCACCTTCCCAAA | 101 |
| *Slc5a2* | F - TGAGTGGAATGCGCTCTTTG<br>R - GAGGCATGGTAATCACTCCG | 86 |
| *Slc9a3* | F - CATGAGCTGAATTTGAAGGATGC<br>R - GCTGAAGTCCACATTGACCAT | 114 |

*Gapdh* - Glyceraldehyde-3-Phosphate Dehydrogenase; *Actb* - Beta-actin; *Ppia* - peptidylproly isomerase A (cyclophilin A); *Slc5a2* - Solute Carrier Family 5 Member 2 (SGLT2); *Slc9a3* - Solute Carrier Family 9 Member A3 (NHE3); F - forward; R - reverse.

### SUPPLEMENTAL FIGURES

#### Figure S1. CONSORT-Style Diagram.

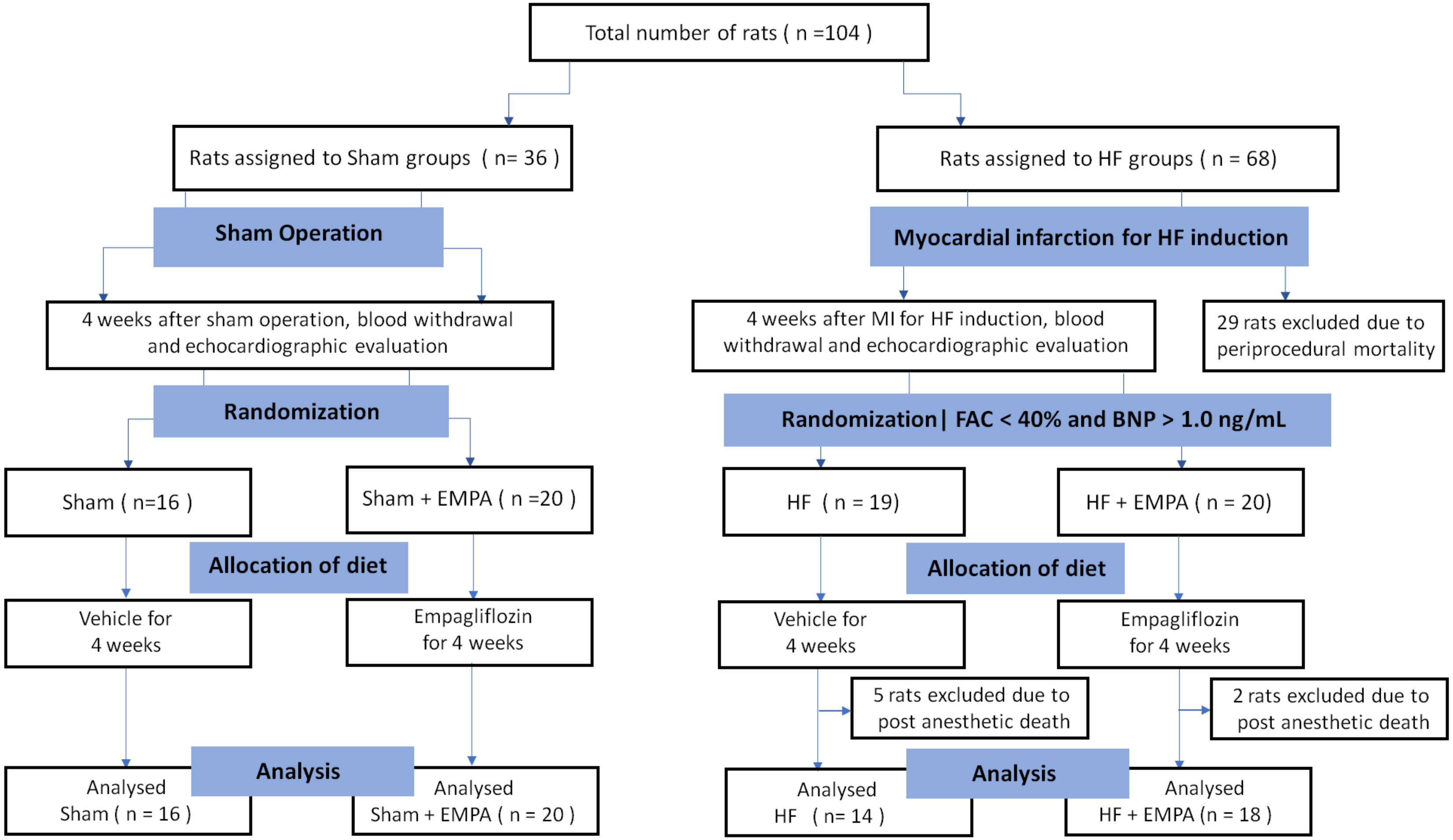

**Figure S2. Schematic timeline of the study**

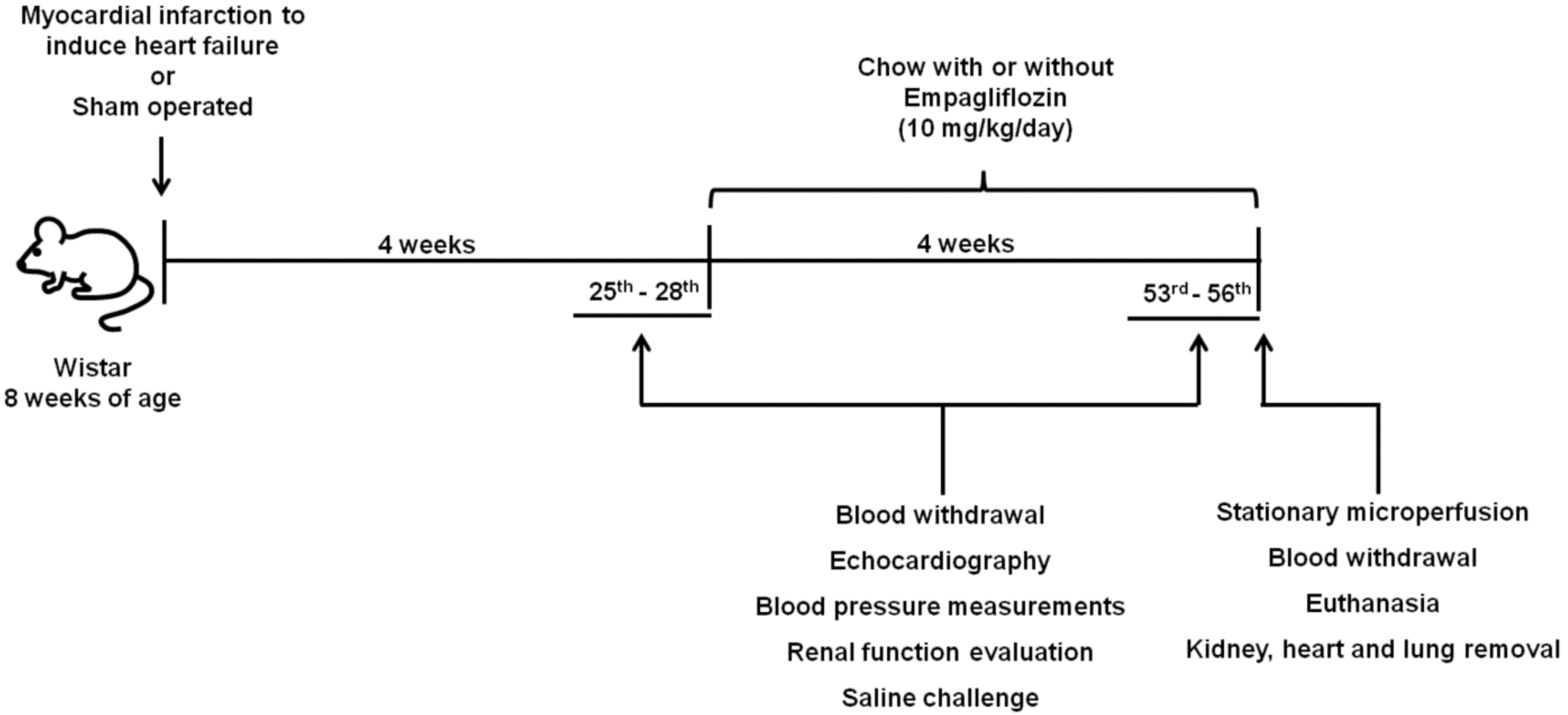

**Figure S3.**

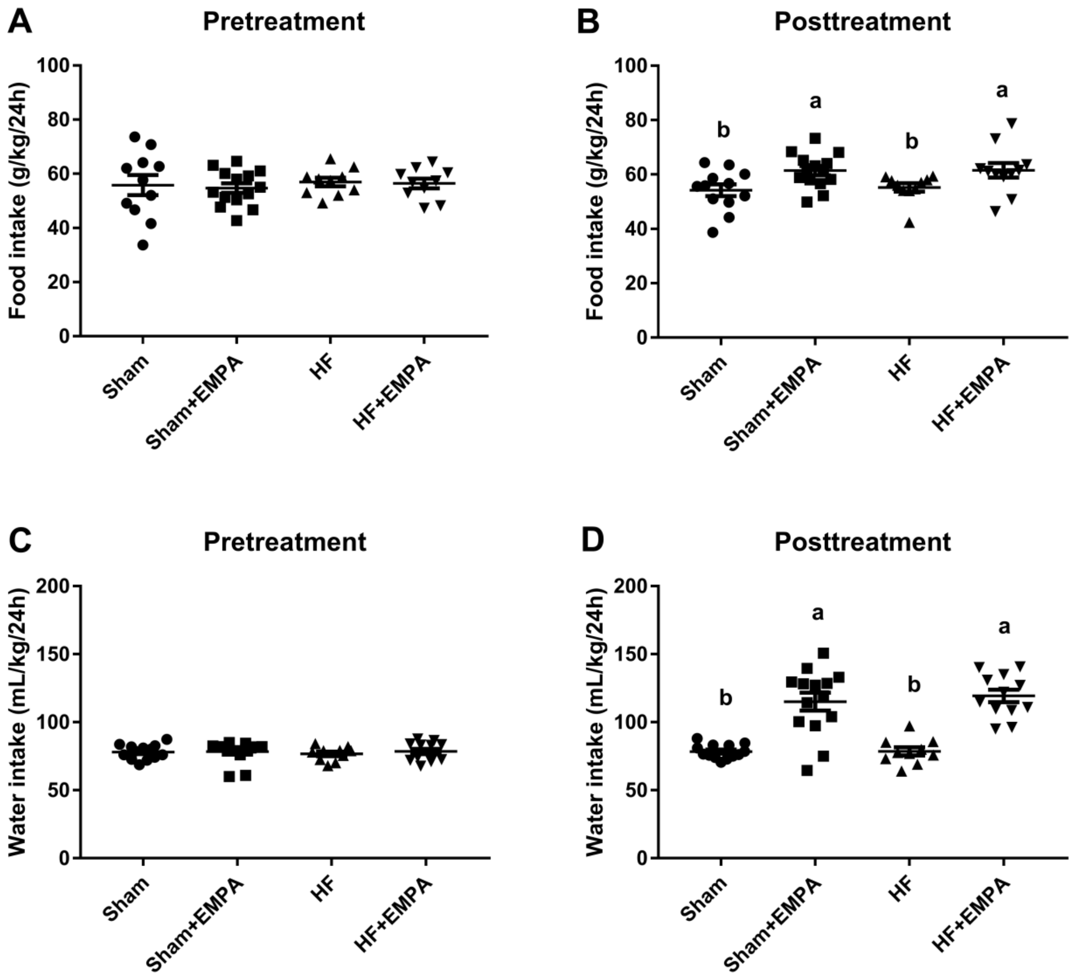


**Figure S3. Evaluation of food and water intake.** Rats were individually placed into metabolic cages for the 24-h urine collection to measure food and water intake. **(A-B)** Food intake. **(C-D)** Water intake. Experiments were conducted four weeks after myocardial infarction for HF induction or sham-surgery (pretreatment) and after treatment with EMPA or not for four weeks (posttreatment). Values represent individual measurements and means ± SEM. Scatter plots with different lowercase letters are significantly different ($P < 0.05$).

**Full unedited gels for Figure 3**

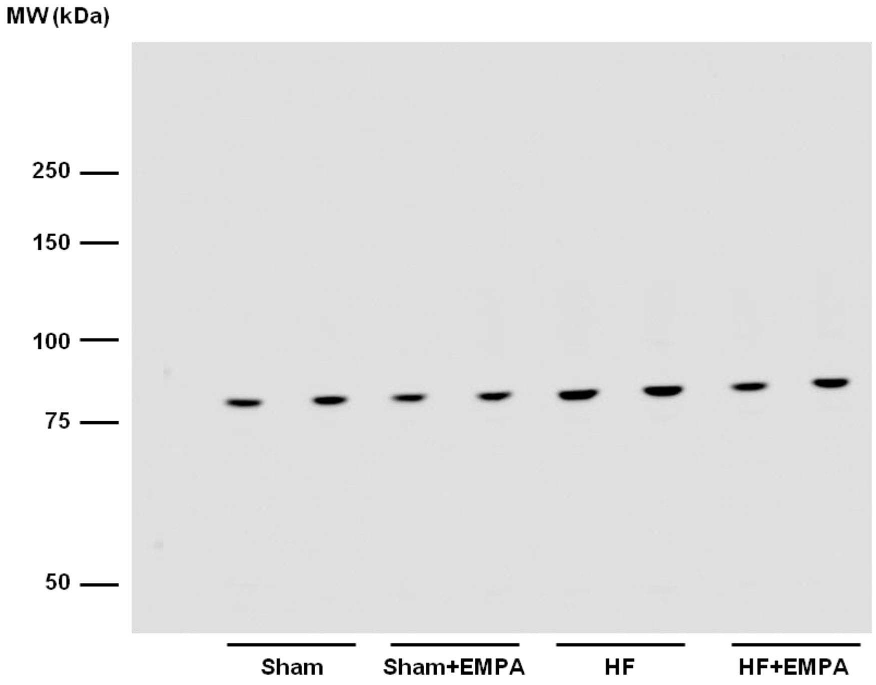


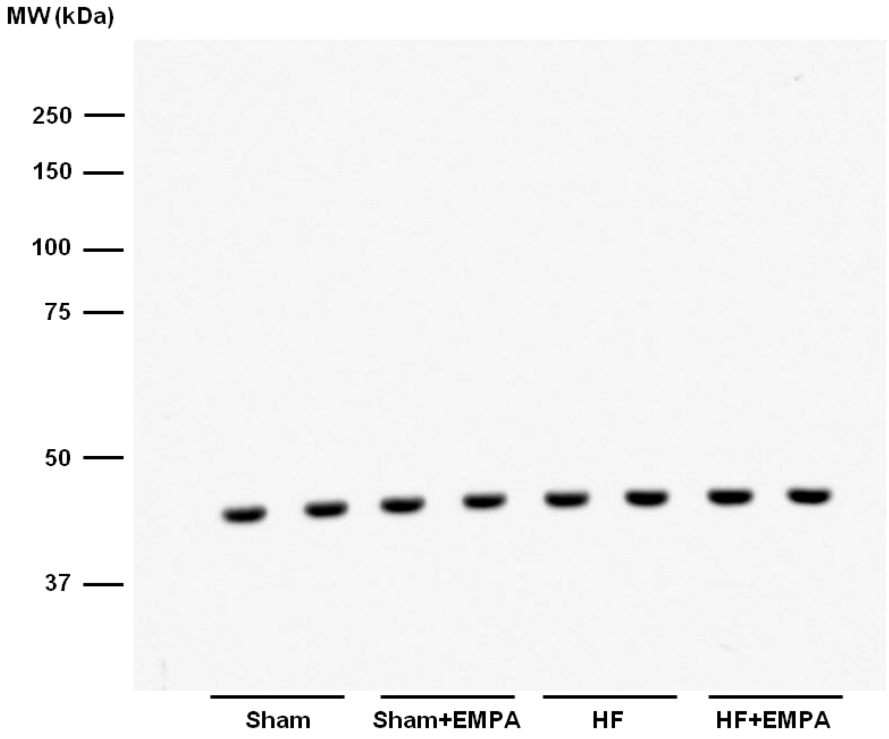

### Full unedited gels for Figure 6

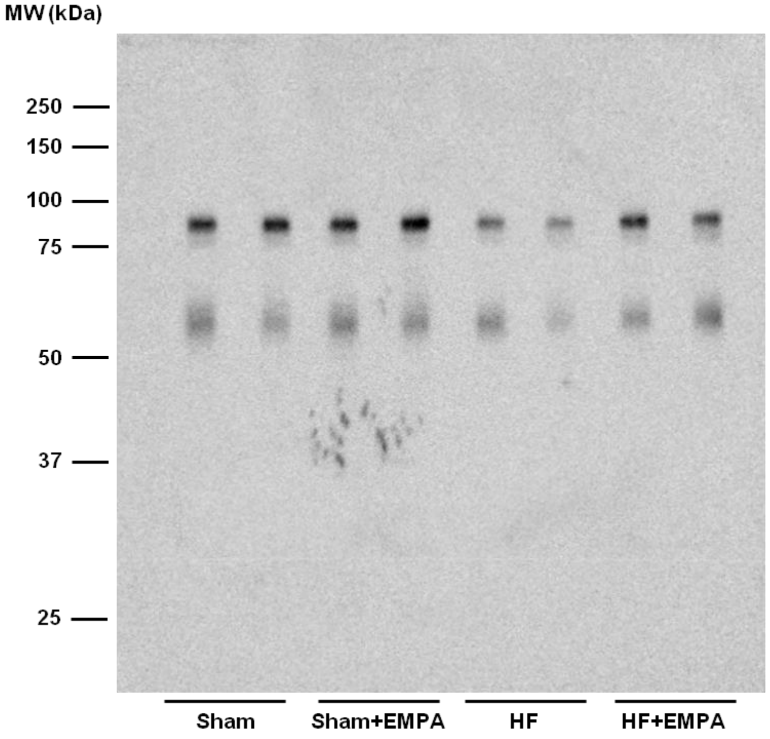


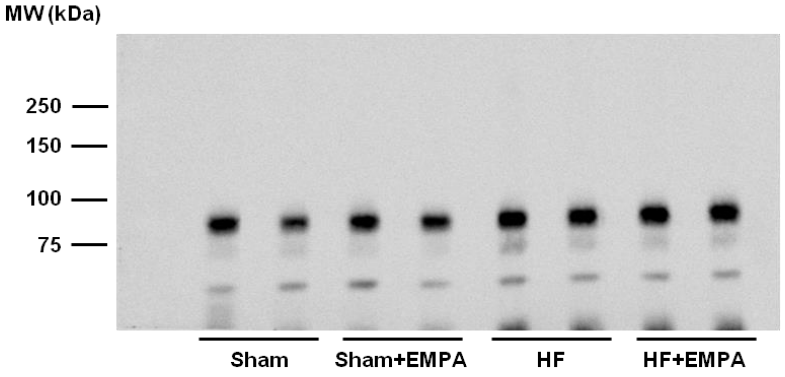


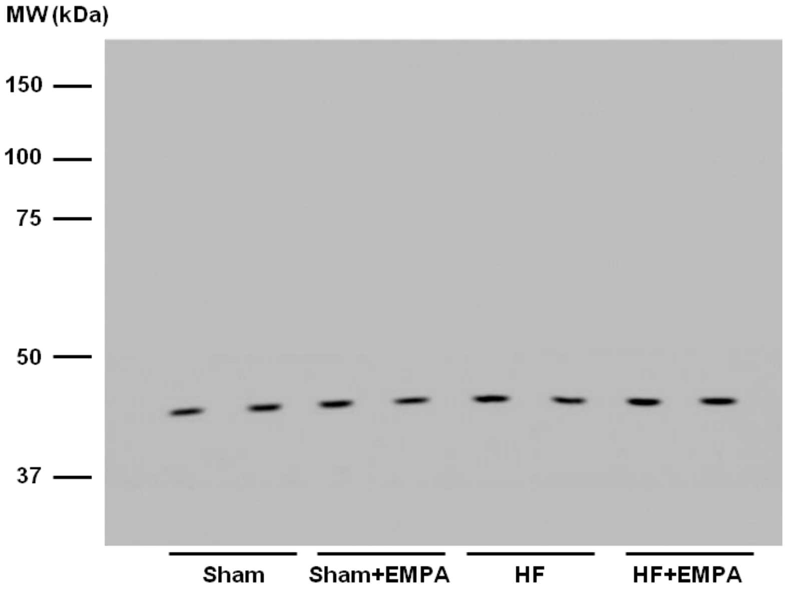
